# Azithromycin and ciprofloxacin have a chloroquine-like effect on respiratory epithelial cells

**DOI:** 10.1101/2020.03.29.008631

**Authors:** Jens F. Poschet, Elizabeth A. Perkett, Graham S. Timmins, Vojo Deretic

**Affiliations:** Department of Molecular Genetics and Microbiology, University of New Mexico Health Science Center; Department of Pediatrics, University of New Mexico Health Science Center; Department of Internal Medicine, University of New Mexico Health Science Center; Department of Pharmaceutical Sciences, University of New Mexico Health Science Center; Department of Autophagy, Inflammation and Metabolism Center of Biomedical Research Excellence, University of New Mexico Health Science Center

## Abstract

There is interest in the use of chloroquine/hydroxychloroquine (CQ/HCQ) and azithromycin (AZT) in COVID-19 therapy. Employing cystic fibrosis respiratory epithelial cells, here we show that drugs AZT and ciprofloxacin (CPX) act as acidotropic lipophilic weak bases and confer in vitro effects on intracellular organelles similar to the effects of CQ. These seemingly disparate FDA-approved antimicrobials display a common property of modulating pH of endosomes and trans-Golgi network. We believe this may in part help understand the potentially beneficial effects of CQ/HCQ and AZT in COVID-19, and that the present considerations of HCQ and AZT for clinical trials should be extended to CPX.

## Introduction

The coronaviruses (CoVs) are a family of viruses causing disease in humans and animals, including the 4 endemic human CoVs (HCoV-NL63, HCoV-OC43, HCoV-229E and HCoV-HKU1) causing mild respiratory tract infections of the common cold category ^1^. The ongoing and raging 2019-2020 pandemic of serious respiratory disease caused by a novel coronavirus ^2,3^ was foreshadowed by two notable prior epidemics: in 2002-2004, severe acute respiratory syndrome (SARS) caused by SARS coronavirus (SARS-CoV; 8,096 infections in 26 countries with 774 deaths) ^4^; and, in 2015, Middle East respiratory syndrome (MERS), caused by MERS coronavirus (MERS-CoV; 2,494 cases in 27 countries with 858 deaths) ^5^, all exhibiting high mortality, with high prevalence of acute respiratory distress syndrome (ARDS), arrhythmia, and death, and, in resolved cases, long-term reduction in lung function and disability. Currently, there are no approved therapeutics or vaccines for coronavirus diseases.

The novel SARS-CoV-like virus, 2019-nCoV (CoV2), related to SARS-CoV but distinct form it, was identified as the causative agent of the current, 2019-2020 outbreak of viral pneumonia that started in Wuhan, China, which spread to all permanently inhabited continents. The human-to-human spread of CoV2 has resulted in a runaway global pandemic, causing a disease termed COVID-19 (<u>c</u>orona<u>v</u>irus disease-20<u>19</u>) by the World Health Organization. COVID-19 pandemics is escalating at an alarming rate with 649,904 infected, as of this writing, and 15,835 deaths just in Italy and Spain with a combined 9.6% mortality rate in these disproportionately affected countries away from the epicenter of the initial epidemic ^6,7^. COVID-19 threatens worldwide populations based on its pernicious combination of long incubation times coupled with exceptional spreading potential, and significantly high morbidity and mortality. The reproductive number R_0_ initially estimated to be 2.2-2.7 ^8,9^ according to some estimates may possibly reach 4.7-6.6 ^10^. There is an urgent public health need to understand CoV2 as a virus, COVID-19 as a disease, and COVID-19 as a frightening epidemiological phenomenon. The world is literally scrambling to develop countermeasures, including therapeutics aiming to lessen disease severity and to come up with prophylaxes including vaccines. Recently, two FDA approved drugs, chloroquine (CQ) and azithromycin (AZT), have shown therapeutic effects in in COVID-19, using viral loads as endpoints in compassionate use/initial clinical trials associated with life-threatening pulmonary complications ^11^.

This work began at the time when a major progress has been made in understanding the key properties of the anomalous ion conductances across cystic fibrosis (CF) respiratory epithelia, with defects in chloride transport via the cystic fibrosis transmembrane conductance regulator CFTR, and in sodium transport via the epithelial sodium channel ENaC ^12^. Certain drugs have proven useful in treating pulmonary disease in CF, such as AZT, an antibiotic that has been a staple of clinical management of CF ^13,14^. CF patients treated with AZT benefit ^14^ in several aspects of lung function, have decreased rate of exacerbations, and experience improved quality of life ^15–19^, albeit the magnitude of these benefits may vary ^20^. As an antibiotic, the mechanism of AZT in CF might be expected to be killing of bacteria, yet the tissue levels of this macrolide do not reach the necessary anti-pseudomonal concentrations in the lungs to provide an easy explanation for its benefits. Whereas several studies propose effects of AZT on bacterial biofilms ^21^, there is compelling evidence that host-mediated effects of this macrolide are just as important since: (i) AZT has beneficial effects in CF, even before ^19^ *Pseudomonas aeruginosa* colonization^22^ occurs; and (ii) AZT protects the bronchial epithelium during *P. aeruginosa* infection independently of its antimicrobial activity ^23^. Thus, AZT confers a previously undefined and antibiotic-independent benefit that is both documented in CF clinical trials ^15–19^ and routinely observed in CF management. However, a subsequent study provided a cautionary note for prolonged AZT’s use in CF since it, like chloroquine, inhibits autophagy ^24^, which is involved in control of mycobacteria ^25^.

Here, we report that AZT and ciprofloxacin (CPX), as has been previously demonstrated for chloroquine ^26^ alter the pH within the intracellular organelles in respiratory epithelial cells, and that this correction results in a normalization of the cell-autonomous immune functions of respiratory epithelia in CF. This effect can be achieved by a diverse set of FDA approved drugs ^26–31^. Our data indicate that AZT’s action at least in part overlaps with CQ’s mode of action. We furthermore propose that clinical trials with patients at risk of developing severe COVID-19, where drugs with deacidifying action such as chloroquine are planned, warrant consideration of CPX,

## Results and Discussion

### Azithromycin corrects organellar pH in CF lung epithelial cells

We tested the effects of AZT on CF respiratory epithelial cells in the absence of bacteria or microbial innate immunity agonists. Among the tests employed, we examine the pH of intracellular organelles since prior studies ^27–29,32–35^ have indicated that these organelles display subtly increased acidification in CF respiratory epithelial cells. Of further relevance, unlike most other macrolides that show little benefit in CF ^14^, AZT possesses two weak base functional groups with pK_a_ of 8.2 and 8.6 (Fig. 1A). The pH of the lumen of the *trans-*Golgi network (TGN) and recycling endosomes (RE) in CF lung cells were measured by ratiometrically applying an intracellular pH-sensitive probe, pHlourin GFP ^36^. The pH sensitive GFP was fused to TGN38 or cellubrevin thus localizing the pHluorin GFP to the lumen of the TGN or RE ^36^ in CF respiratory epithelial cells ^35 34^. Primary CF bronchial epithelial cells treated with AZT displayed correction of the previously reported TGN and RE hyperacidifcation ^27–29,32–35^ (Fig. B-G). Treatment of IB3-1 (CF cells) with either 100 µM (for 1 h) or 1 μM AZT (for 48 h), led to a correction of TGN pH from 6.2 ± 0.1 (control) to 7.1 ± 0.1 or to 6.7 ± 0.2 (Fig. 1C), comparable to the pH of 6.7 ± 0.1 within the TGN of S9 cells (CFTR-corrected IB3-1 cells). In primary CF bronchial epithelial cells (Fig 1D), AZT treatment corrected their TGN pH from 6.1 ± 0.2 to 6.7 ± 0.1 (Fig 1D). Of note is that the absolute concentration of AZT was not critical for the effect at equilibrium, as expected from the extracellular sink of an acidotropic agent accumulating in acidified compartments by being protonated and trapped within the organellar lumen.

**Figure 1.**
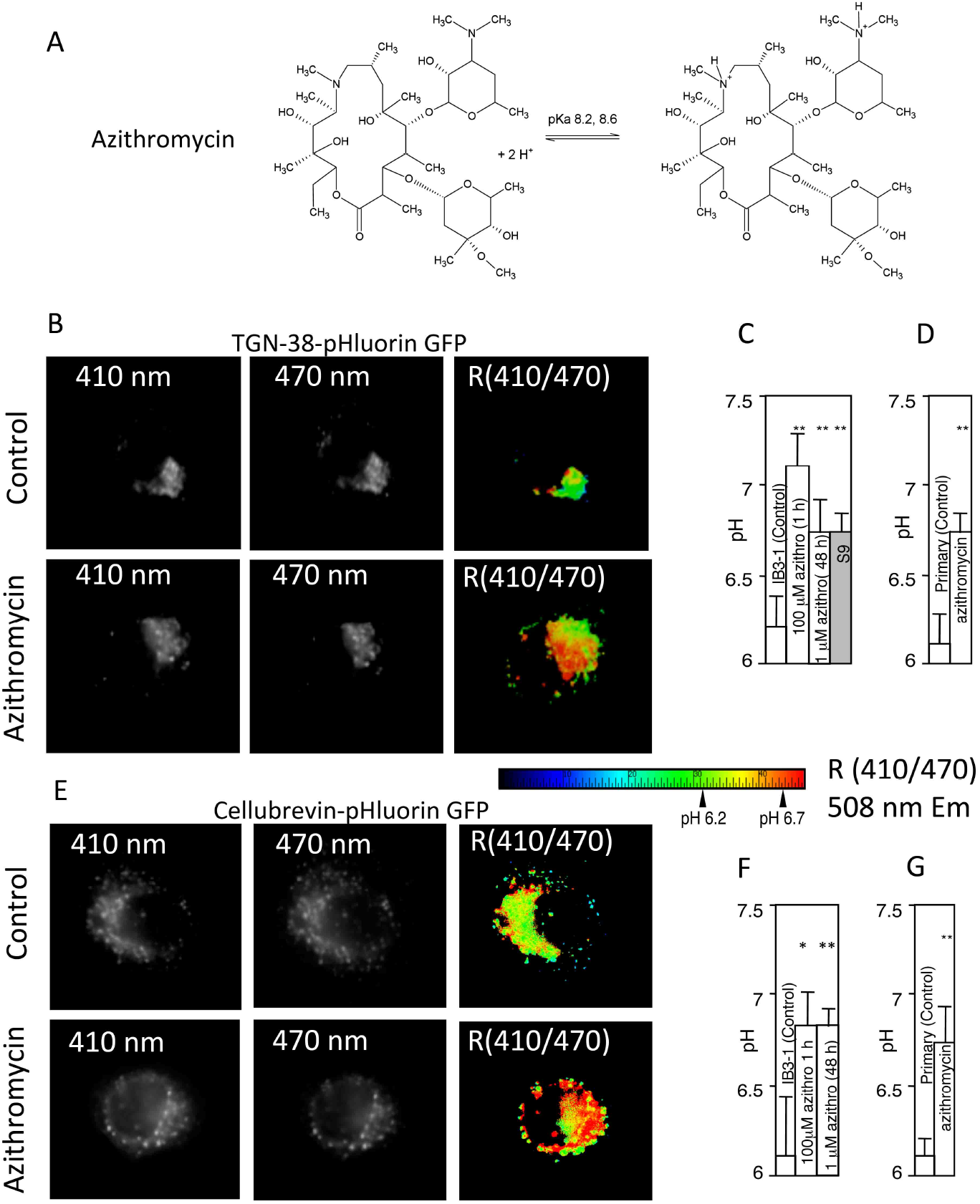
AZT corrects pH in the lumen of intracellular organelles in CF lung epithelial cells. A. AZT chemical formula and pK of its amino groups. B-G. Cystic fibrosis (IB3-1 or primary bronchial epithelial cells from CF lung transplants) and normal (S9, CFTR-corrected IB3-1 cells) were transfected with either TGN38-GFP pHluorin (B-D) or cellubrevin-GFP pHluorin (E-G) and pH of TGN (B-D) or recycling endosome (E-G) determined ratiometrically (emission intensity at 508 nm upon excitation at 410 nm and 470 nm) with or without AZT treatment. Images in B and E: top row, untreated primary bronchial CF cell; bottom row, primary bronchial CF cell treated with AZT. Color look-up table, R (410/470) ratio of emission intensity at 508 nm upon illumination at 410 and 470 nm. C, TGN pH in IB3-1 (CF) cells with or without treatment with AZT, or in S9 (CFTR-corrected IB3-1 cells). D, correction of TGN pH in primary CF respiratory epithelial cells by AZT. E, ratiometric fluorescence images with cellubrevin-GFP pHluorin in primary CF respiratory epithelial cells from lung transplant. F, correction with AZT of cellubrein endosome pH in CF cells (IB3-1). G. Correction with AZT of pH in the cellubrevin endosomes in primary CF respiratory epithelial cells, lung transplant, *P* (ANOVA), * <0.05; ** <0.01, n=6.

The pH of the recycling endosome in CF was similarly corrected with AZT (Fig. 1E-G). Treatment of IB3-1 (CF cells) with 100 µM AZT for 1 h or 1 μM for 48 h led to a correction of RE pH from 6.1 ± 0.3 to 6.8 ± 0.2 (Fig. 1F). In primary CF bronchial cells, 100 µM AZT corrected the pH of RE from 6.1 ± 0.1 to pH 6.7 ± 0.2 (Fig. 1E,G).

### Acidotropic correction of acidified intracellular compartments in CF respiratory epithelial cells is not restricted to azithromycin

CPX, another antibiotic (fluoroquinolone) (Fig. 2) effective in CF is also a weak base with pKa 8.76 ^37^. We tested CPX effects on CF cells. We used pCEP-R cells, the human normal bronchial epithelial cells (16HBE) transfected with R-region of CFTR, which renders them phenotypically express CF characteristics, and found that CPX (100 µM, 1 h) corrected pH in the TGN (Fig. 2). This shows that pH correction in CF is not limited to AZT but that other antibiotics with weak base chemical properties may display this property.

**Figure 2.**
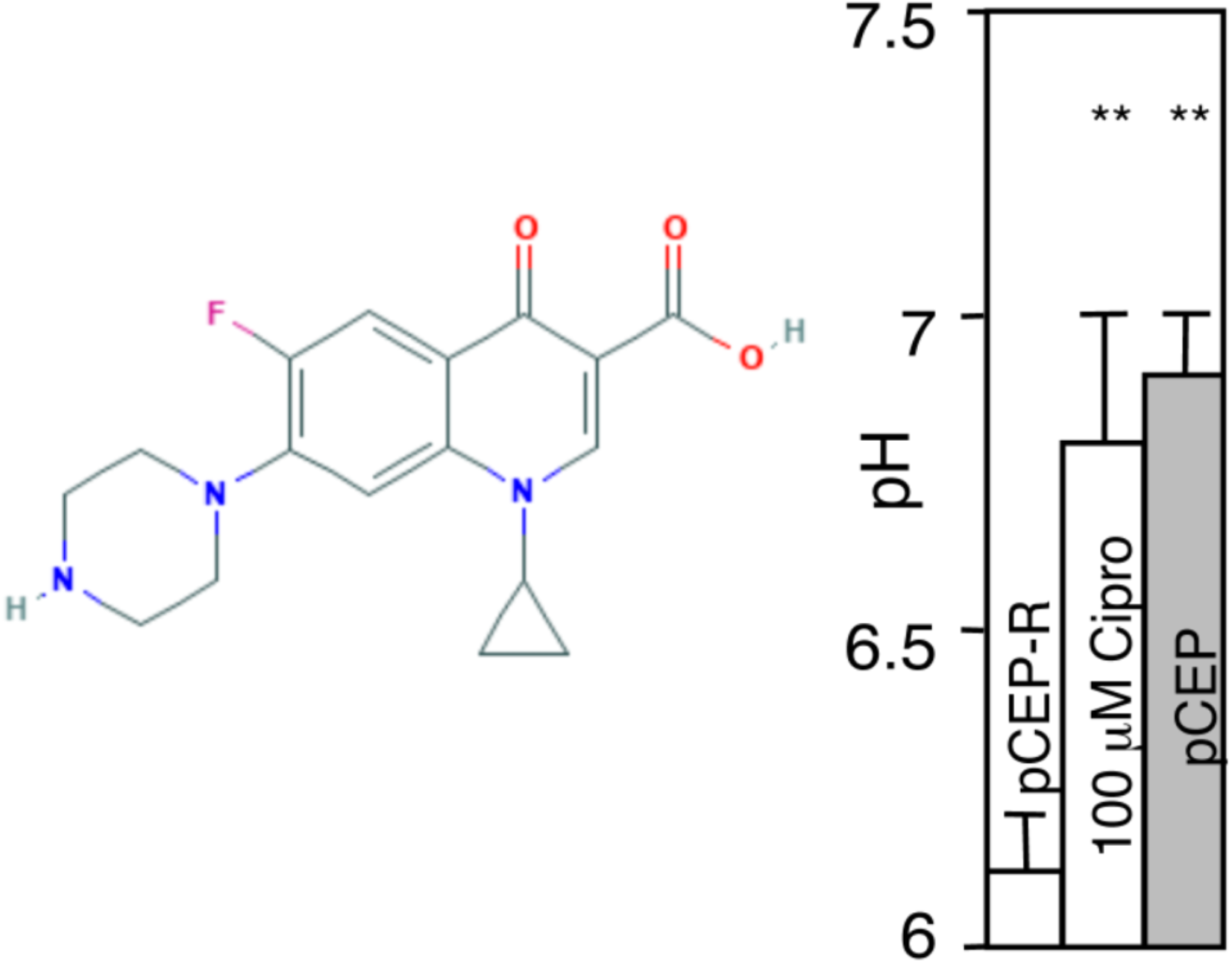
Ciprofolxacin elevates TGN pH in pCEP-R cells. Phenotypically CF cells (pCEP-R) were transfected with TGN38-GFP pHlourin. Transfected cells were treated with 10 µM CPX for 1 h and pH determined ratiometrically as in studies with AZT. pCEP, normal cells.

### Azithromycin corrects cell-autonomous innate immunity-related properties and responses of CF cells

Clinical studies have shown that adherence of *P. aeruginosa* to the epithelia of CF patients undergoing AZT therapy is reduced ^38^. Based on our findings that AZT corrects acidification of TGN in CF cells, we investigated whether AZT affected the undersialylation of aGM1 and reduced bacterial adherence to CF lung epithelial cells. It has been proposed that undersialylation and reduced binding of cholera toxin was one of the evolutionary drivers for the emergence of CFTR mutations in human populations, and that this can be explained at least in part by hyperacidification of the TGN and the resulting suboptimal distribution or activity of sialyltransferases and other glycosylating enzymes in this organelle ^29,39^. CF and normal cells were grown for 14 days post-confluency, followed by 48 h treatment of the monolayers with 1-100 µM AZT. The monolayers were assayed for binding of cholera toxin subunit B (CTB) conjugated to fluorophore FITC (CTB binds preferentially to sialylated glycoconjugates) (Fig. 3A). Using binding of CTB to S9 cells (CFTR-corrected IB3-1 cells) as 100%, and binding to IB3-1 cells (CF) as 0%, treatment of IB3-1 monolayers with 1-100 µM AZT increased CTB-FITC binding to levels that approached that of the non-CF S9 cells (Fig. 3A). As with the TGN-38 pH correction, the effect of AZT on the correction of CTB binding was concentration independent, as expected from accumulation of an acidotropic agent and its intraorganellar trapping once it is protonated.

**Figure 3.**
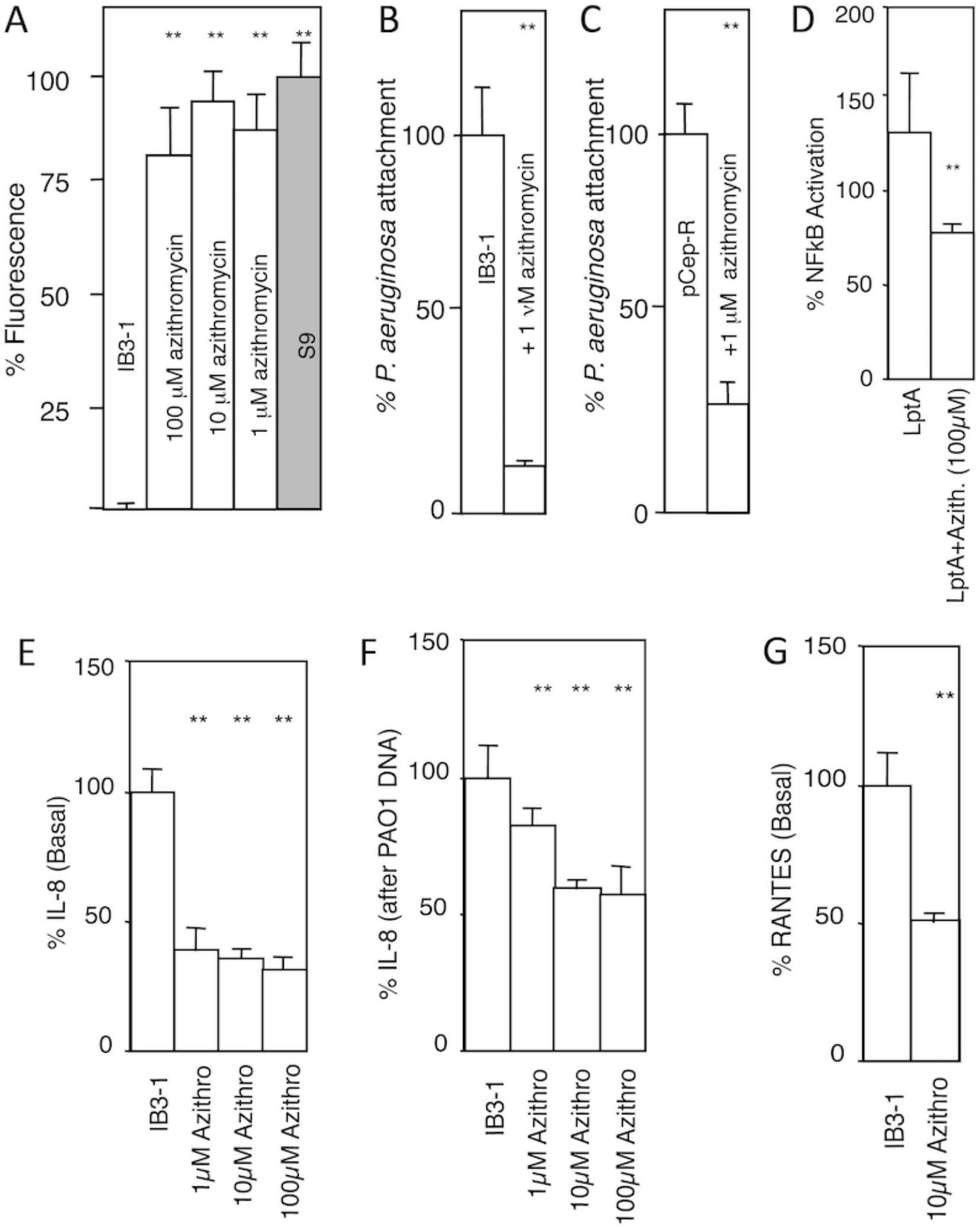
AZT corrects aberrant cell-autonomous innate immunity properties and responses in CF respiratory epithelial cells. A. Cholera toxin (CTB-FITC fluorescence; used as aGM1 probe) binding to CF and normal cell monolayers grown for 14 days post-confluency and treated or not treated with azithomycin (48 h). IB3-1, CF cell line; S9, CFTR-ciorrecetd IB3-1 cells. B and C. *P. aeruginosa* (MOI of 1:200) attachment to IB3-1 (CF) or pCepR-16HBE (normal human bronchial epithelial cells rendered CF by expression of the R domain of CFTR) monolayers pretreated or not treated with AZT (the drug was absent from the adhesion assay). D. AZT reduces NFκB activation (luciferase reporter) in CF cells (IB3-1) in response to *P. aeruginosa* TLR2 ligand derived from lipoprotein LptA. E and F, AZT reduces basal IL-8 secretion and IL-8 induced by treatment with *P. aeruginosa* DNA. G, AZT reduces basal RANTES production by CF (IB3-1) cells. ** *P* (ANOVA; n=6) < 0.05.

We next tested AZT effects on *P. aeruginosa* adhesion to CF cells. We used AZT at a 200-fold lower concentration (1 μM) than its MIC (170 µM) against *P. aeruginosa* ^40^. In addition, the monolayers were washed to eliminate extracellular AZT during bacterial adhesion assay. Pretreatment of IB3-1 (CF) cells with 1 µM AZT led to a decrease in *P. aeruginosa* binding by 85 ± 3% (Fig. 3B). This result was confirmed using pCEP-R cells, the human normal bronchial epithelial cells 16HBE transfected with R-region of CFTR. Pretreatment of pCEP-R (CF) cells with 1 µM AZT led to a decrease of *P. aeruginosa* binding by 75 ± 7% (n = 6, *P* = 0.0002) (Fig. 3C), confirming the findings with IB3-1 CF cells.

### AZT corrects immune response of CF cells to challenges with bacterial products

We tested the effects of AZT on the well-accepted pro-inflammatory phenotype of the CF respiratory epithelial cells ^41–44^, by measuring NFκB, TGF-β, IL-8, and RANTES responses. When IB3-1 cells were treated with 100 µM AZT, the basal NFκB activation was reduced by 50 ± 8 % compared to untreated cells. When CF-cells (IB3-1) were stimulated with *P. aeruginosa* lipopeptide (LptA) for 6 h in the presence of 100 µM AZT, the macrolide attenuated the increased NFκB stimulation in response to LptA by 40% (Fig. 3D). These findings are in keeping with reports that AZT ^42^ and weak bases (chloroquine) or inhibitors (bafilomycin A_1_) of the vacuolar H^+^ ATPase ^45^ ameliorate proinflammatory signaling in CF cells.

The IL-8 levels are increased in CF ^41,43,44,46–48^, and this IL-8 increase can be corrected by treatment with weak bases such as chloroquine ^45^. We tested the effect of AZT on IL-8 release under basal conditions and upon stimulation with *P. aeruginosa* DNA. Treatment of CF-cells with 1 to 100 µM AZT reduced basal IL-8 secretion by almost 70 % (Fig. 3E). Stimulation of IL-8 production by addition of *P. aeruginosa* DNA led to a 5-fold increase of IL-8 secretion (unstimulated = 55 pg IL-8 / 100 µg protein vs. DNA stimulated = 240 pg IL-8 / 100 µg protein). Pretreatment with AZT of stimulated cells reduced IL-8 secretion (10 µM AZT 33 ± 7%, 10 µM AZT 34 ± 10%,) (Fig. 3F). As with the other assays, concentration of AZT in the medium was not critical to obtain effects although longer pre-incubation times (e.g. 24-48 h for 1 μM AZT vs. 1-4 h for 100 μM AZT) were needed at lower concentrations. It is important to note that this behavior is in support of the AZT action as a weak base that accumulates in acidified intracellular organelles, which act as a sink for trapping AZT upon its protonation. In addition, the effect of AZT on basal RANTES levels ^45^ was measured and AZT reduced RANTES secretion (Fig. 3G).

### AZT corrects furin activity and TGF-β levels in CF cells

Genetic polymorphisms ^49^ associated with elevated profibrotic mediator TGF-β are key disease modifiers in CF, with elevated TGF-β increasing disease severity ^50–53^. We have reported that TGF-β generated by CF respiratory epithelial cells is increased compared to normal cells and that treatment with the weak base CQ can correct TGF-β release ^27,33^. When AZT was tested, it too reduced TGF-β (Fig. 4A). We have reported that the elevated TGF-β in CF cells is caused by increased furin activity in the intracellular organelles of CF respiratory epithelial cells ^27^. Hence, we tested whether AZT affected the furin inbalance. When CF cells were incubated with 100 µM AZT, production of active furin was normalized in CF cells, on par with CQ treatment and on level with the CFTR-corrected S9 cells (Fig. 4B). These results indicate that correcting organellar pH with AZT normalizes the excessive processing and activation of furin in CF cells. In summary, AZT benefits in CF that cannot be explained by its direct antibiotic activity are due to its physical-chemical action as a weak base, normalizing organellar function and cell-autonomous responses of CF respiratory epithelial cells.

**Figure 4.**
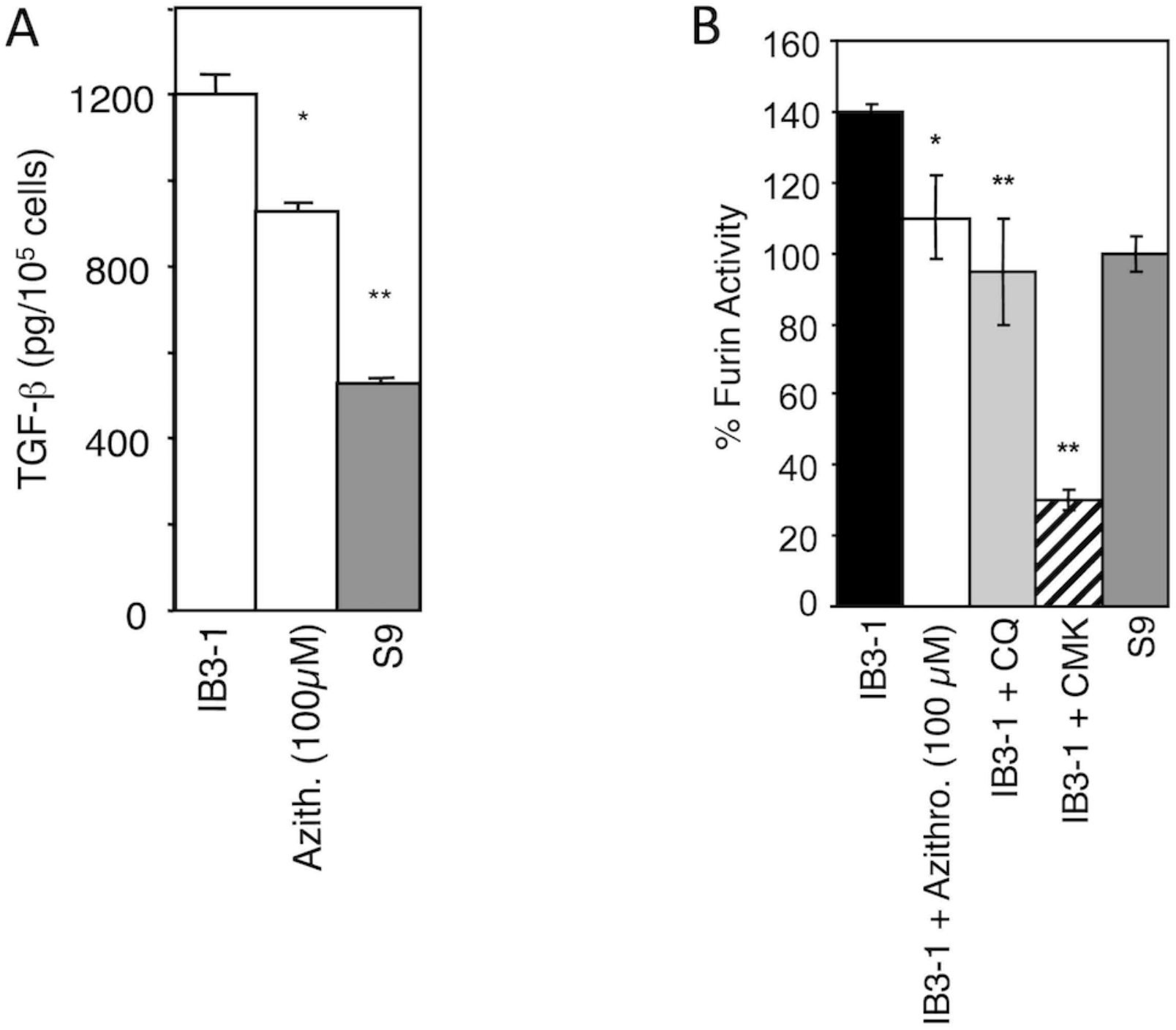
AZT reduces TGF-β production and furin activity in CF respiratory epithelial cells. A. AZT effect on TGF-β secretion by CF (IB3-1) cells. B. Furin levels were corrected in full medium with AZT or 0.1 mM chloroquine (CQ). 50 µM furin inhibitor CMK was used as a measure of maximum furin inhibition. * *P* < 0.01, ** *P* < 0.05 (ANOVA, n=6).

### Pharmacological consideration of relevance for COVID-19

The selected properties of the alternate drugs AZT and CPX are shown in Table 1, together with those of CQ and HCQ. All are hydrophobic weak base drugs, with amine pKa from 8.2 to 10.1, in accordance with their hypothesized common mechanism. The amount of base absorbed per 100 mg of compound is also calculated, and when combined with the typical dosage ranges and schedules, shows that the overall likely amounts of base for intracellular acid neutralization from each are similar in amount for HCQ, AZT and CPX.

**Table 1.**
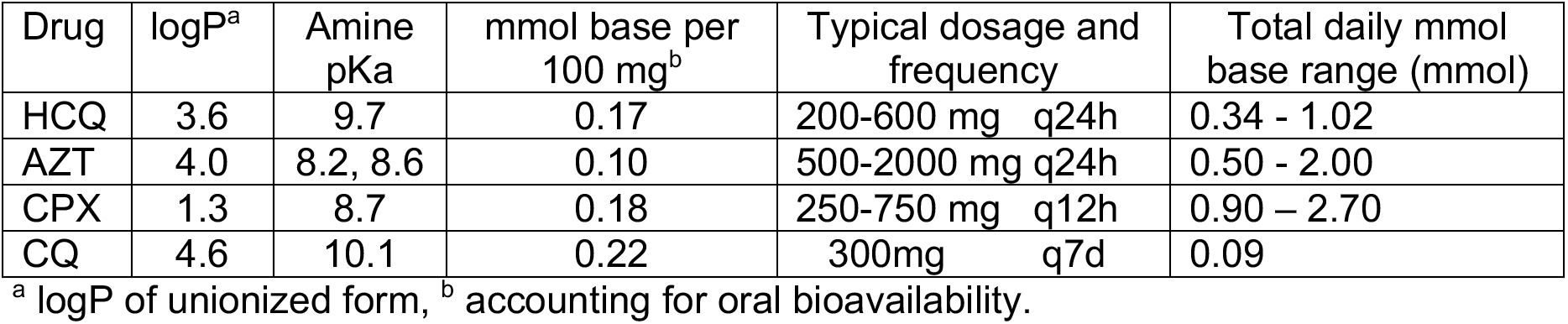
Selected properties of study drugs.

### Conclusions and perspective

In conclusion, the relationships demonstrated in this study indicate that a set of FDA-approved drugs, CQ, AZT, CPX, with known safety profiles, act by controlling pH of intracellular organelles in respiratory epithelial cells, a highly relevant target for CoV2. Furthermore, our previous studies of phosphodiesterase 5 inhibitiors show similar effects, and these may also be further tested. ^30^

The alterations of pH in organelles of the secretory pathway, especially TGN ^31^, may alter glycosylation of both the receptors for CoV2, such as ACE2, which binds the spike protein, and glycosylation of CoV2 proteins during their biogenesis and assembly. Importantly, coronavirus M proteins govern viral budding at the ER-to-Golgi intermediate compartment but penetrate deeper into the Golgi apparatus ^54^, and among the pathogenic CoVs, at least the MERS CoV M protein possesses TGN localization signals ^55^. The effects of chloroquine have been observed with other viruses including HIV gp120 glycosylation defect and loss of infectivity ^56,57^ as well as HSV arrest of noninfectious parties in the TGN ^58^. Our studies here show that CF respiratory epithelial cell normalization is based on a neutralization of the pre-existing pathologically hyperacidified compartments due to the underlying CF disease, resulting in normalization of glycosylation in CF respiratory epithelial cells. We postulate here that the acidotropic and neutralization effects of AZT, CPX and CQ in normal cells would result in vectorially similar pH shifts away from optimal for the function of enzymes within the secretory pathway that would also alter glycosylation.

The alterations of pH in organelles of the endosomal pathway may have effects on host proteases acting upon the CoV2 Spike protein to execute its cleavage necessary for virus entry. The inhibition of viral entry in recent cellular systems is best achieved with a combination of camostat mesylate (an inhibitor of TMPRSS2 protease) plus E-64d, an inhibitor of lysosomal cathepsins ^59^. It has been noted that CoV2, unlike SARS-CoV, possesses a novel furin cleavage site within its Spike protein ^60^. Our studies here show that AZT can reduce furin activity that is abundantly present in respiratory epithelial cells. Multiple proteases are proposed to cleave CoV2 Spike, including TMPRSS2 ^59^ (which also acts on SARS-CoV ^61–63^), furin ^60^, and lysosomal hydrolases ^59^. This warrants consideration of a more general approach to their inhibition, such as altering pH, rather than focusing solely on a specific host protease. We propose that the drugs studied here, CQ, AZT and CPX, may modulate these aspects of host-pathogen interactions and virus infectious cycle, with immediate compassionate use in the clinic or in controlled clinical trials. A validation of the effects of CPX in COVID-19 patients would enable the utilization of this drug to alleviate likely shortages in HCQ and AZT. CPX is a part of the US Health and Human Services Strategic National Stockpile ^64^, which can be mobilized during regional or national emergencies.

## Methods

### Cells, tissue culture and chemicals

IB3-1 is a bronchial epithelial cell line derived from a CF patient with a ΔF508/W1282X *CFTR* mutant genotype ^65^. S9 is a derivative of IB3-1 corrected for chloride conductance by stable transfection with a functional *CFTR*. The cells were maintained in LHC-8 Media (Biosource Int., MD). The CF cell line CFBE41ō (ΔF508/ΔF508) and normal human bronchial epithelial cells16HBEō (from D. Gruenert) were maintained in complete MEM-media (GibcoBRL, Life Technologies, MD). The normal human tracheal cell line 9HTEō, transfected with pCEP-R or pCEP were from P. Davis, and were maintained in complete DMEM (GibcoBRL, Life Technologies, MD). Primary bronchial epithelial cells were from Clonetics (NHBE) or from P. Karp (CF lung transplants) and were maintained in supplemented BEMB (Lonza, MD). AZT was from Fluke (Switzerland) and CPX-HCL was from Sigma (St. Louis, MO).

### Transfections and microscopy

TGN38 and cellubrevin fusion with pH sensitive GFP was from J. Rothman ^36^. Cells were transected with 1 µg/ml DNA using Effectene (Siegen, CA) for 48 h prior to measuring pH. Co focal fluorescence microscopy of fixed samples was carried out on Zeus 510 META microscope.

### Ratiometric pH determination

The ratio of emission at 508 nm upon excitation at 410 nm vs. 470 nm was obtained described previously ^28,34–36^. A pH standard curve was generated by collapsing the pH-gradient by incubating cells in 10 µM monensin and 10 µM nigericin for 30 min at 37° in buffer at pH 7.4, 6.5 or pH 5.5 and ratios were recorded for internal standards. Fluorescence images were taken upon excitation at 410 and 470 nm (6 consecutive exposures). Three regions of interest were selected and the standard curve was plotted as averaged 410/470 ratio values for a given buffer pH. Cells were incubated with 1-100 µM AZT for 1 h or 48 hrs, prior to determining pH. The ratio 410/470 of the samples was read against the standard curve and converted into pH units.

### Cholera toxin adsorption to CF lung epithelial cells, *P. aeruginosa* adherence to CF respiratory epithelial cells, NF-κB activation, IL-8 and RANTES measurements

CF and normal cells, grown to post confluency, were incubated for 48 h with 1-100 µM AZT. Cells were fixed and stained with fluorescently labeled cholera toxin B for 1 h at room temperature. Relative fluorescence was calculated from actual grey levels of 10 random fields from 3 independent experiments. Postconfluent cells, treated with 1-100 µM AZT, were incubated with *P. aeruginosa* (PAO1) at a multiplicity of infection of 1:200 in MEM media without serum or antibiotic supplements. Samples were washed 3 times with PBS and lyses with 0.5% TX-100. Cell associated bacteria were plated on *P. aeruginosa* isolation agar and colonies counted. Treatment with *Pseudomonas* pro-inflammatory lipopeptide (lipotoxin) Lepta, NF-κB activation (luciferase assay), IL-8 and RANTES analysis were carried out as previously described ^45,66^.

TGF-β production and furin activity analysis. TGF-β levels were determined using the PAI/L assay as described previously ^67^. Furin enzyme activity was assayed in the presence of 0.25% Triton X-100 as a permeabilizing agent using boc-RVRR-amc (100 µM) as a furin substrate. Fluorescence was measured at 380 nm excitation and 460 nm emission wavelengths. The inhibitory concentration of dec-RVKR-cmk (Calbiochem) for ExoA-mediated cell death, were assessed and 50 µM was employed in subsequent experiments.

Statistics. Unless otherwise specified, statistical analyses were carried out using Fisher’s Protected LSD post hoc test (ANOVA) (SuperANOVA, Abacus Concepts).

## Acknowledgments

We thank P. Karp and J. Zabner for primary bronchial epithelial cells from CF patient lungs, P. Davis and P. Zeitlin for cell lines. This work was supported by NIH grants AI 50825, AI 31139 and AI81015, and Cystic Fibrosis Foundation (CFF) Grants DERETI08G0 and TIMMIN03I0. JFP was a CFF postdoctoral fellows. VD was supported in part by the NIH grants R37AI042999, R01AI042999 and a center grant P20GM121176.

